# Unexpected implications of STAT3 acetylation revealed by genetic encoding of acetyl-lysine

**DOI:** 10.1101/537696

**Authors:** Yael Belo, Zachery Mielko, Hila Nudelman, Ariel Afek, Oshrit Ben-David, Anat Shahar, Raz Zarivach, Raluca Gordan, Eyal Arbely

**Affiliations:** Department of Chemistry, Ben-Gurion University of the Negev, Beer-Sheva, 8410501, Israel; Center for Genomic and Computational Biology, Department of Biostatistics and Bioinformatics, Duke University, Durham, NC 27708, USA; The National Institute for Biotechnology in the Negev, Ben-Gurion University of the Negev, Beer-Sheva, 8410501, Israel; Department of Life Sciences, Ben-Gurion University of the Negev, Beer-Sheva, 8410501, Israel; Department of Computer Science, Department of Molecular Genetics and Microbiology, Duke University, Durham, NC 27708, USA

**Author notes:** Corresponding author *Email address* (Eyal Arbely).

**Keywords:** Lysine acetylation, deacetylation, protein-DNA interaction, post-translational modifications crosstalk, genetic code expansion

## Abstract

The signal transducer and activator of transcription 3 (STAT3) protein is activated by phosphorylation of a specific tyrosine residue (Tyr705) in response to various extracellular signals. STAT3 activity was also found to be regulated by acetylation of Lys685. However, the molecular mechanism by which Lys685 acetylation affects the transcriptional activity of STAT3 remains elusive. By genetically encoding the co-translational incorporation of acetyl-lysine into position Lys685 and co-expression of STAT3 with the Elk receptor tyrosine kinase, we were able to characterize site-specifically acetylated, and simultaneously acetylated and phosphorylated STAT3. We measured the effect of acetylation on the crystal structure, and DNA binding affinity and specificity of Tyr705-phosphorylated and non-phosphorylated STAT3. In addition, we monitored the deacetylation of acetylated Lys685 by reconstituting the mammalian enzymatic deacetylation reaction in live bacteria. Surprisingly, we found that acetylation, *per se*, had no effect on the crystal structure, and DNA binding affinity or specificity of STAT3, implying that the previously observed acetylation-dependent transcriptional activity of STAT3 involves an additional cellular component. In addition, we discovered that Tyr705-phosphorylation protects Lys685 from deacetylation in bacteria, providing a new possible explanation for the observed correlation between STAT3 activity and Lys685 acetylation.

## 1. Introduction

Signal transducer and activator of transcription 3 (STAT3) is a member of the STAT protein family of latent transcription factors that are activated in response to the binding of cytokines, growth factors and hormones to extracellular receptors [1, 2]. Structurally, STAT3 comprises an N-terminal domain, followed by a coiled-coil domain, a DNA-binding domain, a Src homolodgy 2 (SH2) domain and a C-terminal transactivation domain [3–5]. According to the canonical Janus kinase (JAK)-STAT pathway, receptor tyrosine-phosphorylation, promoted by the binding of signaling molecules to cell surface receptors, is followed by SH2 domain-mediated binding of STAT3, which is then phosphorylated on Tyr705. Tyrosine phosphorylation enables STAT3 dimerization by reciprocal binding between the SH2 domain of one STAT3 monomer and the phosphorylated Tyr705 (pY705) of the other STAT3 monomer. Dimeric STAT3 then accumulates at the nucleus, where it acts as a transcription factor by binding to specific DNA response elements [6]. That being said, it was found that non-phosphorylated STAT3 is transcriptionally active, and that STAT3 may serve non-canonical roles, such as, for example, in mitochondria [5, 7–11].

STAT3 was found to be constitutively activated in various cancer cell lines and tumor tissues, where it promotes tumor cell proliferation, invasion, and migration [12, 13]. As a transcription factor that mediates extracellular signaling and gene transcription, STAT3 is also involved in the communication between cancer cells and the microenvironment [14]. In addition, STAT3 plays metabolic, developmental and anti-inflammatory roles [15]. These diverse activities of STAT3 are regulated by various mechanisms, including an array of post-translational modifications, such as phosphorylation (e.g., Tyr705), methylation and acetylation. Specifically, acetylation of Lys685 was suggested to be important for STAT3 dimerization and full transcriptional activity [16, 17]. Lys685 acetylation was also found to promote STAT3-DNA methyltransferase 1 interactions, and subsequent methylation of tumor-suppressor promoters [18]. However, the exact biochemical mechanism of Lys685 acetylation-dependent transactivation is not fully understood. According to crystal structures of STAT3, the relatively flexible side-chain of Lys685 is not directly involved in mediating dimerization [3, 5]. Therefore, it is not clear if and how the intra-dimer interface is affected by Lys685 acetylation. In addition, Stark and co-workers found that acetylation of Lys685 is important for the transcriptional activity of unphosphorylated, but not of phosphorylated, STAT3 [19]. Furthermore, acetylation of Lys685 was suggested to increase in response to cytokine-mediated stimulation [17, 20], although Chen and co-workers found that CD44 can mediate Lys685 acetylation in a cytokine- and growth factor-independent manner [21].

## 2. Materials and Methods

### 2.1. General

General chemicals and DNA oligomers for molecular cloning were ordered from Sigma Aldrich (Darmstadt, Germany). DNA sequencing was performed by the sequencing facility at Ben-Gurion University. The STAT3 gene was amplified from pDONR221 plasmid carrying STAT3 cDNA (DNASU plasmid ID HsCD00295594). pBK vector for expression of evolved acetyl-lysine synthetase was kindly provided by Dr. Jason W. Chin (MRC-LMB, Cambridge, UK).[22] Enzymes for molecular cloning were purchased from NEB (Ipswich, MA) and used according to the manufacturer’s instructions. DNA was purified using spin columns from Macherey Nagel (Düren, Germany). Acetyl lysine was purchased from Chem-Impex International Inc. (Wood Dale, IL) and used without further purification. DH10B *E. coli* strain (Life technologies, Carlsbad, CA) was used for molecular cloning and plasmid propagation. BL21(DE3) *E. coli* strain (NEB, Ipswich, MA) was used for protein expression. Bacteria were incubated in liquid LB media or on LB/agar plates supplemented with antibiotics (50 *μ*g/mL kanamycin, spectinomycin, or chloramphenicol). Primary antibodies: anti-6×His (#G020) was purchased from abm (Richmond, ON); anti Y705-phosphorylated STAT3 (#ab76315) was purchased from Abcam (Cambridge, UK); anti K685-acetylated STAT3 (#PA5-17429) was purchased from Thermo Fisher Scientific (Waltham, MA). Secondary antibodies: anti-mouse IgG (#ab7068) and anti-rabbit IgG (#ab92080) were purchased from Abcam.

### 2.2. Molecular cloning

Non-phosphorylated STAT3 (residues 128–715, accession number NP_644805.1) was expressed as a fusion protein with C-terminal tobacco etch virus (TEV) cleavage site, followed by the lipoyl domain and 6× His-tag. The gene was cloned into the first open reading frame of a pCDF-Duet vector using Gibson Assembly Kit (NEB, Ipswich, MA).[23, 24] This vector also contained a U25C mutant of PylT under constitutive expression.[25] To enable the co-translational incorporation of acetylated lysine, an in-frame TAG mutation was introduced at position Lys685 by site-directed mutagenesis. To express phosphorylated STAT3 proteins, STAT3 and K685-TAG STAT3 (residues 128–715) with C-terminal 6× His-tag were cloned into the first open reading frame of the above mentioned pCDF-Duet vector. Phosphorylated STAT3 variants were expressed without C-terminal TEV cleavage site and lipoyl domain, since TEV protease was incapable of digesting the Tyr705-phosphorylated variants of STAT3. Next, the gene coding for the kinse domain of the Elk receptor was amplified from TKB1 cells (Agilent Technologies, Santa Clara, CA) and cloned with a C-terminal HA-tag between NdeI and EcoRV restriction sites within the second open reading frame of the pCDF-Duet vector. Thus, this plasmid enabled the IPTG-controlled co-expression of STAT3 (either acetylated or non-acetylated) and Elk.

### 2.3. Protein expression

For expression of Lys685-acetylated STAT3, *E. coli* BL21(DE3) cells were co-transformed with a pBK vector for constitutive expression of evolved acetyl-lysine synthetase and the appropriate pCDF vector for expression of either non-phosphorylated or phosphorylated STAT3. Transformed cells were incubated overnight in 2×TY medium, supplemented with 50 *μ*g/mL spectinomycin and 50 *μ*g/mL kanamycin. The next day, overnight cultures were diluted to OD_600_ =0.02 into 4 L of 2×TY medium supplemented with the same antibiotics. The cultures were incubated at 37°C until OD_600_ =0.3, when they were supplemented with 10 mM acetylated lysine and 20 mM nicotinamide. At OD_600_ =0.6, protein expression was induced with 0.5 mM IPTG, and the incubation temperature was reduced to 18°C. After 18 h, the cells were harvested by centrifugation, and the pellet was stored at - 80°C. Expression of non-acetylated STAT3 was performed in a similar manner, except that cells were only transformed with the pCDF vector for expression of STAT3 (or co-expression of STAT3 and Elk), and the bacteria were cultured in media supplemented with 50 *μ*g/mL spectinomycin without nicotinamide or acetylated lysine.

### 2.4. Protein Purification

Frozen bacterial pellet (~20 gr) was resuspended in 100 mL of buffer A (50 mM Tris pH 8.0, 100 mM NaCl, 15 mM β-mercaptoethanol, 20 mM imidazole pH 8.0) supplemented with 1.2 *μ*g/mL leupeptin, 1 *μM* pepstatin A, 100 *μM* PMSF, 1 *μ*g/mL aprotinin, 0.4 mg/mL lysozyme, 20 *μ*g/mL DNAse, 10 mM MgCl_2_, and 10 mM nicotinamide. For the purification of Tyr705-phosphorylated STAT3, 100 *μ*M of sodium orthovanadate were add to the buffer. Cells were incubated on ice with stirring for 30 min, lysed by sonication and the lysate was centrifuged at 20,000 g for 30 min at 4°C. Clear lysate was loaded on a 5 mL HisTrap HP column (GE Healthcare, Chicago, IL) pre-equilibrated with buffer A. Column was washed with at least 10 column volumes of buffer A and protein was eluted with a linear gradient (0–100% over 20 column volumes) of buffer B (50 mM Tris pH 8.0, 100 mM NaCl, 15 mM β-mercaptoethanol, 500 mM imidazole pH 8.0). Fractions were analyzed by sodium dodecyl sulfate polyacrylamide gel electrophoresis (SDS-PAGE), and fractions of highest purity were collected. The combined fractions of non-phosphorylated STAT3 variants were diluted 1:3 with dialysis buffer (25 mM Tris pH 7.6, 300 mM NaCl, 10% Glycerol, 15 mM β-mercaptoethanol) and filtered. TEV protease was added to the sample and dialysis was performed against 2×4 L of dialysis buffer at 4°C. Dialyzed protein solution was supplemented with imidazole (25 mM) and the protein was further purified by a second Ni^2+^ affinity chromatography. Flow-through containing the cleaved protein was collected and diluted 1:10 in ice-cold buffer C (25 mM Tris pH 7.6, 10% Glycerol, 15 mM β-mercaptoethanol). When phosphorylated STAT3 variants were purified, dialysis was performed without TEV protease, and dialyzed sample was diluted 1:10 into ice-cold buffer C, without a second Ni^2+^ affinity chromatography. Diluted protein samples in buffer C were loaded on a heparin HP 5 mL column (GE Healthcare, Chicago, IL) and protein was eluted with buffer D (25mM Tris pH 7.6, 10% Glycerol, 15 mM β-mercaptoethanol, 1 M NaCl) using reverse flow. Eluted protein was concentrated using Amicon Ultra-15 centrifugal filter unit with nominal molecular weight limit of 10 kDa (Merck Millipore, Burlington, MA), loaded on a HiLoad 26/600 Superdex 200 column (GE Healthcare, Chicago, IL) pre-equilibrated with buffer E (HEPES pH 7.0, 200 mM NaCl, 5 mM DTT, 10 mM MgCl_2_ and 0.5 mM PMSF) and protein was eluted at a flow rate of 2.6 ml/min with fractionation. Fractions were analyzed by 12% SDS-polyacrylamide gel electrophoresis and fractions of highest purity were combined, concentrated to ~0.5 mg/mL (based on UV absorption, ε=89840) using Amicon Ultra-15 centrifugal filter unit and stored at −80°C.

### 2.5. Fluorescence polarization assay

A DNA probe (5’-AGCAGTTCTGGGAAATCT-3’) modified by fluorescein at the 3’ end was purchased from Integrated DNA Technologies (Coralville, IA). Fluorescein-labeled single-stranded DNA was mixed with the non-labeled complement sequence at a 1:1.1 ratio in phosphate-buffered saline (PBS). The solution was heated to 90°C and the DNA was annealed by a slow temperature gradient (1°C/min). The solution of 3’-fluorescein-labeled double-stranded probe was filtered (0.2 *μ*m) and stored in small aliquots at a final concentration of 20 *μ*M. Fluorescence polarization signals were recorded on a Spark multimode microplate reader (Tecan, Männedorf, Switzerland) in a 384-well format. Aliquots (50 *μ*l) were prepared by mixing 2 nM DNA probe and the indicated STAT3 variant at increasing concentrations, starting at 0.5 *μ*M for Tyr705-phosphorylated variants or 5 *μ*M for non-phosphorylated variants. For each variant, 40 samples were prepared with a 1.3-fold increase in protein concentration between samples, in PBS supplemented with 1 mM DTT, 1 mg/ml bovine serum albumin and 50 *μ*g/ml salmon sperm DNA (R&D Systems, Minneapolis, MN). Fluorescence polarization data were fitted to Equation 1:

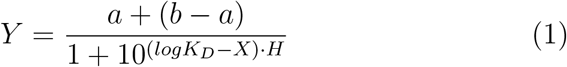

where *Y* is the fluorescence polarization signal, *X* is the STAT3 concentration, *a* and *b* are the minimal and maximal fluorescence polarization signals, respectively, *K_D_* is the dissociation constant, and *H* is the Hill slope.

### 2.6. In vitro DNA binding specificity

In vitro DNA-binding specificities were measured using the standard PBM protocol [26]. All possible 10-mer double-stranded DNA oligonucleotides were first obtained by primer extension. Next, the microarray was incubated 1 h with blocking solution (2% milk in PBS), followed by a 1 h incubation with PBS-based protein-binding solution of 6×His-tagged purified STAT3, and washing step. Bound STAT3 proteins were then labeled with Alexa488-conjugated anti-6 × His antibodies (1 hr incubation, 1:20 dilution, Qiagen 35310). The array was washed and scanned using a GenePix 4400A scanner (Molecular Devices) at 2.5 *μ*m resolution. Data were normalized as previously described [26].

### 2.7. Crystallization

Single-stranded DNA sequences (5’-TGCATTTCCCGTAAATCT-3’ and 5’-AAGA TTTACGGGAAATGC-3’, IDT, Coralville, IA) were dissolve in annealing buffer (1 M Tris-HCl pH 8, 1 M NaCl and 0.5 M EDTA pH 8), heated to 90°C, and annealed by a slow temperature gradient (−1°C/min). Double-stranded molecules were dialyzed extensively against DDW (18.2 MΩ·cm) at 4°C, filtered through a 0.2 *μ*M filter, lyophilized and redissolved in ultra pure water at concentration of 0.175 mM. Purified STAT3 and double-stranded DNA were mixed at 1:2 molar ratio (STAT3 dimer:double stranded DNA), and the sample was incubated on ice for 1 h. Crystals were grown at 20°C using the sitting drop vapor diffusion technique. Drops composed of 2 *μ*l protein/DNA complex and 0.5 *μ*l of crystallization solution were equilibrated above a reservoir of 80 *μ*l. Initial crystallization experiments were performed using the Hampton Research index screen (Aliso Viejo, CA). Final crystallization conditions were 0.1 M Bis-Tris pH 5.5, and 0.1 M magnesium sulfate for the AcK685 STAT3, and 0.2 M lithium sulfate, 0.1 M Bis-Tris pH 7.0 and 20% w/v polyethylene glycol 3350 for AcK685+pY705 STAT3. Before data collection, crystals were transferred into a cryo-protectant solution consisting of 60% mother liquor, 25% ethylene glycol, and 15% DDW. The protected crystals were mounted on Hampton Research CryoCapHT nylon loops and flash-frozen in liquid nitrogen.

### 2.8. Data collection, structure determination, and data analysis

Diffraction data were collected at the European Synchrotron Radiation Facility (Grenoble, France), beamline ID-29. Data were indexed and integrated with XDS. Initial phase determination was performed by molecular replacement with Phaser from the CCP4 package, using a previously solved STAT3 structure (PDB: 1BG1)[27] as the search model. The structure was further refined using CCP4 Phenix.[28] Successive rounds of model building and manual corrections were performed with COOT.[29] Figures were prepared using PyMol.

### 2.9. Deacetylation assay in bacteria

Deacetylation assay was performed as previously described [30]. Briefly, *E. coli* ΔCobB BL21(DE3) cells were transformed with pBK vector for constitutive expression of an evolved acetyl-lysine synthetase. The transformed bacteria were made competent and co-transformed with a pACYC-Duet vector encoding one of the human deacetylases (Sirt1–7 and HDAC6_479–835_) and a pCDF vector for the expression of AcK685 STAT3, or AcK685+pY705 STAT3. Cells were recovered in 1 mL SOC (37°C, 600 rpm) and incubated for 16-18 h (37°C, 220 rpm) in LB media supplemented with 50 *μ*g/mL kanamycin, spectinomycin, and chloramphenicol. Overnight cultures were diluted to OD_600_=0.05 into 5 mL of pre-warmed (37°C) auto-induction medium [31], supplemented with 10 mM AcK and 25 *μ*g/mL chloramphenicol and spectinomycin each, and 50 *μ*g/mL kanamycin. Cultures were incubated at 37°C (220 rpm) for 6 h, when the temperature was lowered to 22°C for an additional 42 h. Western blot analysis was performed using antibodies against the C-terminal 6× His tag and K685-acetylated STAT3.

### 2.10. Western blot

Protein-expressing bacteria from 1 mL of overnight culture normalized to O.D_600_=1 were precipitated (15,000 rpm, 10 min, 4°C) and cell pellet was resuspended in 500 *μ*L of 1× Laemmli sample buffer. Cells were lysed by heating to 95°C with agitation (400 rpm) for 7 min and cell debris were precipitated by centrifugation at 4°C (15,000 rpm). Proteins in equal volumes of cleared lysates were separated by SDS-PAGE and transferred to a 0.2 *μ*m nitrocellulose membrane using a semi-dry transfer apparatus (Trans-Blot Turbo, BioRad, Hercules, CA). Membranes were blocked with Tris-buffered saline containing 0.05% (v/v) Tween-20 (TBST) and 5% (w/v) non-fat dry milk, and incubated over night with primary antibody diluted in 5% (w/v) bovine serum albumin in TBST, at 4°C. On the following day, membranes were washed with 1× TBST, incubated with secondary antibody for 1 h at room temperature, and washed again. Finally, proteins were visualized using ECL reagent (GE Healthcare, Chicago, IL) and immunoblot intensities were quantified with ImageJ.[32].

### 2.11. MS/MS

Purified protein samples were separated by SDS-PAGE, and the band corresponding to ~64 kDa was incised and in-gel digested by trypsin according to the manufacturer’s protocol (Promega). Peptides were then extracted from the gel and analyzed by LC-MS using an Eksigent nano-HPLC (model nanoLC-2D, Netherlands) connected to an LTQ Orbitrap XL ETD mass spectrometer (Thermo Fisher Scientific, Germany & USA). Reverse-phase chromatography of peptides was performed using a C-18 column (IntegraFrit, 360 μm OD × 75 μm ID; New Objective USA). Peptides were separated by a 70 min linear gradient, starting with 100% buffer A (5% acetonitrile, 0.1% formic acid) and ending with 80% buffer B (80% acetonitrile, 0.1% formic acid), at a flow rate of 300 nl/min. A full scan, acquired at 60,000 resolution, was followed by CID MS/MS analysis performed for the five most abundant peaks, in the data-dependent mode. Fragmentation (with minimum signal trigger threshold set at 500) and detection of fragments were carried out in the linear ion trap. Maximum ion fill time settings were 500 ms for the high-resolution full scan in the Orbitrap analyzer and 200 ms for MS/MS analysis in the ion trap. The AGC settings were 5 × 10^5^ and 1 × 10^4^ (MS/MS) for Orbitrap and linear ion trap analyzers, respectively. Proteins were identified and validated using the SEQUEST and Mascot search engines operated under the Proteome Discoverer 1.4 software (Thermo Fisher Scientific). Mass tolerance for precursors and fragmentations was set to 10 ppm and 0.8 Da, respectively. Only proteins containing at least two peptides of high confidence (Xcore 2 or 2.5 or more for doubly or triply charged species, respectively) were chosen.

## 3. Results

### 3.1. Expression of AcK685 and AcK685+pY705 STAT3

A widely accepted experimental approach in functional studies of lysine acetylation involves the use of Lys-to-Arg or Lys-to-Gln mutants. In a different experimental approach, acetylation levels are modified by deacetylase inhibitors, or by knockdown or over-expression of acetyltransferases or histone deacetylases. However, the former approach may have unknown structural and functional effects, while the latter may indirectly affect cell physiology. Therefore, we decided to study site-specifically acetylated STAT3 by genetically encoding the co-translational incorporation of Nε-acetyl lysine in response to an in-frame TAG stop codon at position 685 of the STAT3 core domain (residues 128-715) [22, 33–36]. Lys685-acetylated STAT3 was produced by co-expression of K685-TAG *stat3* with pyrrolysine amber suppressor tRNA (PylT) and evolved acetyl lysine synthetase (AcKRS) [22]. The Tyr705-phosphorylated protein was obtained by co-expression of STAT3 with the protein-tyrosine kinase domain of the Elk receptor, as previously demonstrated [27, 37]. To express STAT3 site-specifically modified by acetylation and phosphorylation, K685-TAG *stat3* was co-expressed with the amber suppression machinery for co-translational incorporation of an acetylated lysine residue at position Lys685, together with the kinase domain of Elk receptor for post-translational phosphorylation of Tyr705.

The incorporation of acetyl-lysine at position 685 was validated by Western blot analysis using specific antibodies against Lys685-acetylated STAT3 (Supplementary Figure S1 A), and trypsin digestion followed by tandem MS/MS (Supplementary Figure S2). Similarly, Tyr705-phosphorylation was dependent on co-expression with the Elk kinase domain, and was validated by immunoblotting using specific antibodies against Tyr705-phosphorylated STAT3 (Supplementary Figure S1B) and MS/MS (Supplementary Figure S2). This experimental setup enabled the expression of four STAT3 variants, namely wild type (WT), Lys685-acetylated (AcK685), Tyr705-phosphorylated (pY705), and Lys685-acetylated+Tyr705+phosphorylated (AcK685+pY705) STAT3. Subsequent comparative studies of WT and AcK685 STAT3 should, therefore, report on effects of acetylation on non-phosphorylated STAT3, while studies comparing pY705 and pY705+AcK685 should report on effects of Lys685 acetylation on phosphorylated STAT3.

### 3.2. Lys685-acetylation-dependent in vitro DNA-binding affinity and specificity

The effect of Lys685 acetylation on in vitro DNA-binding affinity was measured by fluorescence anisotropy. A 3’-fluorescein-labeled oligonucleotide derived from the binding site of the α2-macroglobulin (α2M) promoter (5’-AGCAGTT-CTGGGAAATCT-3’) was incubated with increasing concentrations of the four STAT3 variants, and the fluorescence polarization signal as a function of STAT3 concentration was fitted to Equation 1. Similar *K_D_* values were measured for pY705 STAT3 and AcK685+pY705 STAT3 (25±2 nM and 20±3 nM, respectively), demonstrating that Lys685 acetylation had little to no effect on the in vitro affinity of Tyr705-phosphorylated STAT3 to the binding site of the α2M promoter (Figure 1). Similar results were obtained when measurements were performed in the presence of 0.1 or 1 mg/mL nuclear protein extract from STAT3 null PC3 cells (Supplementary Figure S3). In addition, the *K_D_* values of WT STAT3 and AcK685 STAT3 were higher than 1000 nM, suggesting that non-phosphorylated STAT3 had low affinity for the binding site of the α2M promoter, with no significant effect of Lys685 acetylation on DNA-binding affinity. Therefore, our data show that the addition of an acetyl group to Lys685, *per se*, had no effect on STAT3 DNA-binding affinity.

**Figure 1:**
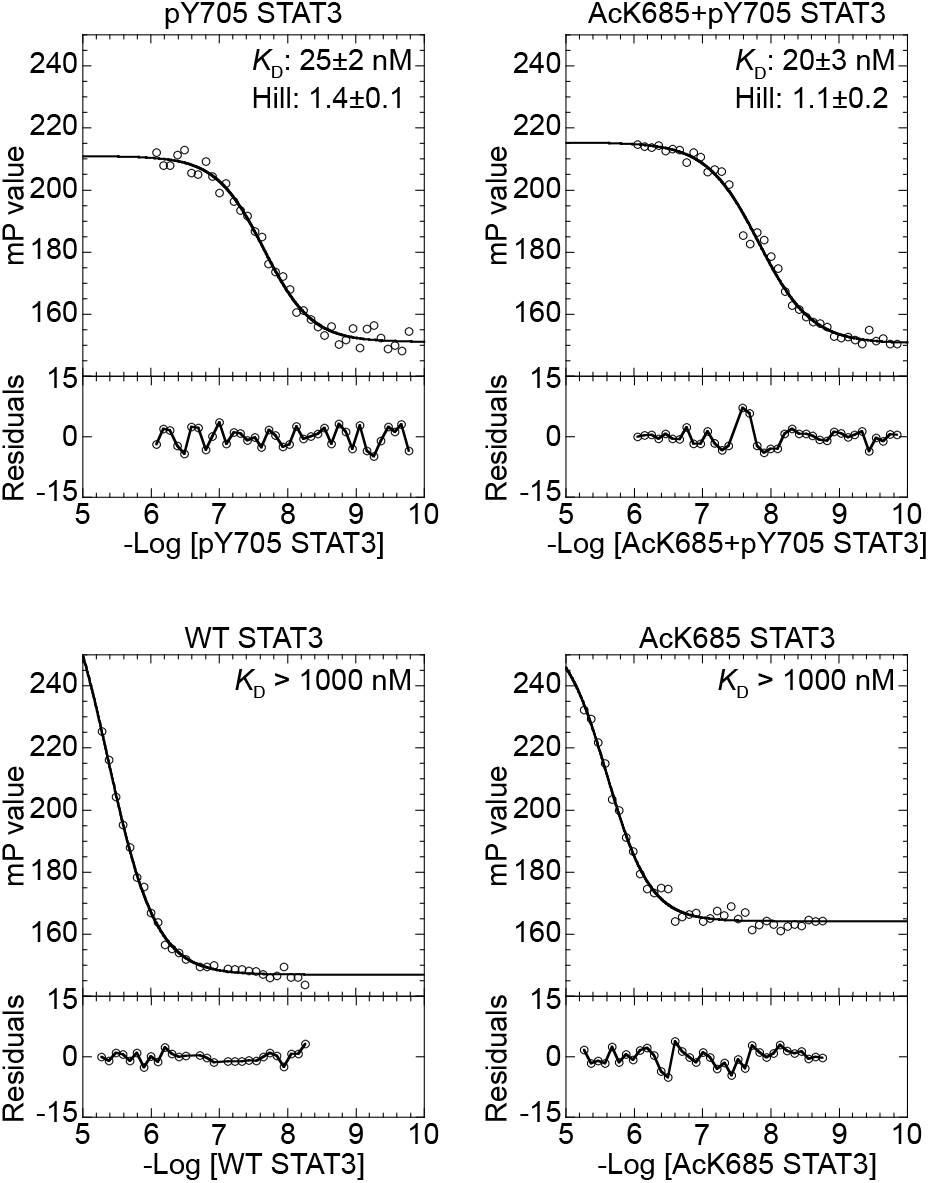
Affinity of STAT3 variants to the α2M promoter, measured by fluorescence anisotropy. Increasing concentrations of the indicated STAT3 proteins were incubated with 2 nM of 3’-fluorescein-labeled probe, and data were fitted to Equation 1. Average *K_D_* and Hill slope values are displayed (*n*=3, ±SD). The affinity of non-phosphorylated STAT3 variants was below the detection limit. mP: millipolarization units.

It has been demonstrated that post-translational modifications can affect the DNA-binding specificity of transcription factors, and consequently, their transcriptional activity [24]. Therefore, we asked whether Lys685 acetylation affected the DNA-binding specificity of Tyr705 phosphorylated STAT3. To answer this question, we comprehensively characterized the DNA-binding specificities of pY705 STAT3 and AcK685+pY705 STAT3 in an unbiased manner, using the well-established protein-binding microarray (PBM) technology (Figure 2) [26]. Binding of STAT3 to all possible 10-mer nucleotides was detected and position weight matrix (PWM) logos, the most common way to represent protein-DNA binding specificity [39], were then generated for the two STAT3 variants using the most enriched DNA sequences (Figure 2). We found that both pY705 STAT3 and AcK685+pY705 STAT3 had the highest affinity to the same 9 nucleotide-long sequence (5’-TTCC(G/C)GGAA-3’), with essentially indistinguishable PWM logos. These PWM logos strongly suggest that acetylation of Lys685 has no effect on the in vitro DNA-binding specificity of Tyr705-phosphorylated STAT3.

**Figure 2:**
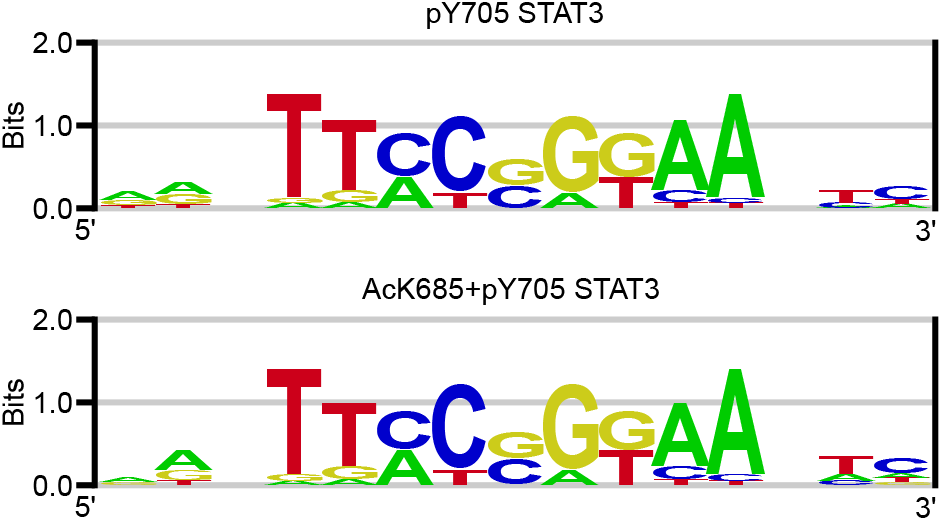
In vitro STAT3 DNA-binding specificity. Position weight matrix logos for pY705 STAT3 and AcK685+pY705 STAT3, obtained from a universal protein-binding microarray containing all possible 10 mer combinations on a single chip [38]. The measurements provide two indistinguishable PWM logos, suggesting an identical DNA-binding specificity for pY705 STAT3 and AcK685+pY705 STAT3.

### 3.3. Crystal structure of Lys685-acetylated and Tyr705-phosphorylated STAT3 in a complex with DNA

To study the effect of Lys685 acetylation on protein-protein and protein-DNA interactions, we co-crystallized AcK685+pY705 STAT3 with double-stranded DNA (5’-AAGATTTACGGGAAATGC-3’). The complex of AcK685+ pY705 STAT3 with DNA was crystallized in the *P*4_1_ space group and the structure was solved to a resolution of 2.85 Å (PDB ID: 6QHD; statistics of data collection and model refinement are listed in Supplementary Table S1). The asymmetric unit was composed of a STAT3 dimer bound to a double-stranded DNA molecule, in contrast to other STAT3 crystal structures, which contain only one STAT3 monomer with single-stranded DNA within the asymmetric unit [3, 5].

Within each monomer, electron density map for most of our protein model was well defined, yet as with the crystal structure of pY705 STAT3 (PDB ID: 1BG1)[3], several residues, including loops within the SH2 domain, were poorly defined and consequently were not included in the final model; i.e., the loop connecting α-helices 1 and 2 (185–193), residues 419–427 between β-sheets e and f, a loop at the end of α-helix 7 (536– 538), and several residues within the SH2 domain (626–632, 658–665, 689–702; all numbers refer to monomer A). Nevertheless, the overall SH2 domain backbone could be traced, and the relative orientation of the SH2 domains was highly similar to that found in the crystal structure of non-acetylated STAT3 (Figure 3A). Moreover, the overall crystal structure of AcK685+pY705 STAT3 in complex with DNA was essentially identical to the structure of pY705 STAT3 (PDB ID: 1BG1) [3]. Superposition of these two structures revealed a C_α_ root mean square deviation (RMSD) value of 0.47 Å, indicative of the high similarity between them, with only minor conformational changes (Figure 3A). In addition, according to the electron density map, the positions of AcK685 and pY705 backbone atoms were not affected by Lys685-acetylation (Figure 3B and C). We also determined the low resolution (>3 A) crystal structure of AcK685 STAT3 in complex with DNA. Superposition of the backbone structures of AcK685+pY705 and AcK685 STAT3 (C_α_ RMSD=0.64 A) revealed that the two crystal structures are essentially identical (Supplementary Figure S4). Taken together, we found no significant effect of Lys685 acetylation on the crystal structure of Tyr705-phosphorylated STAT3 in complex with DNA.

**Figure 3:**
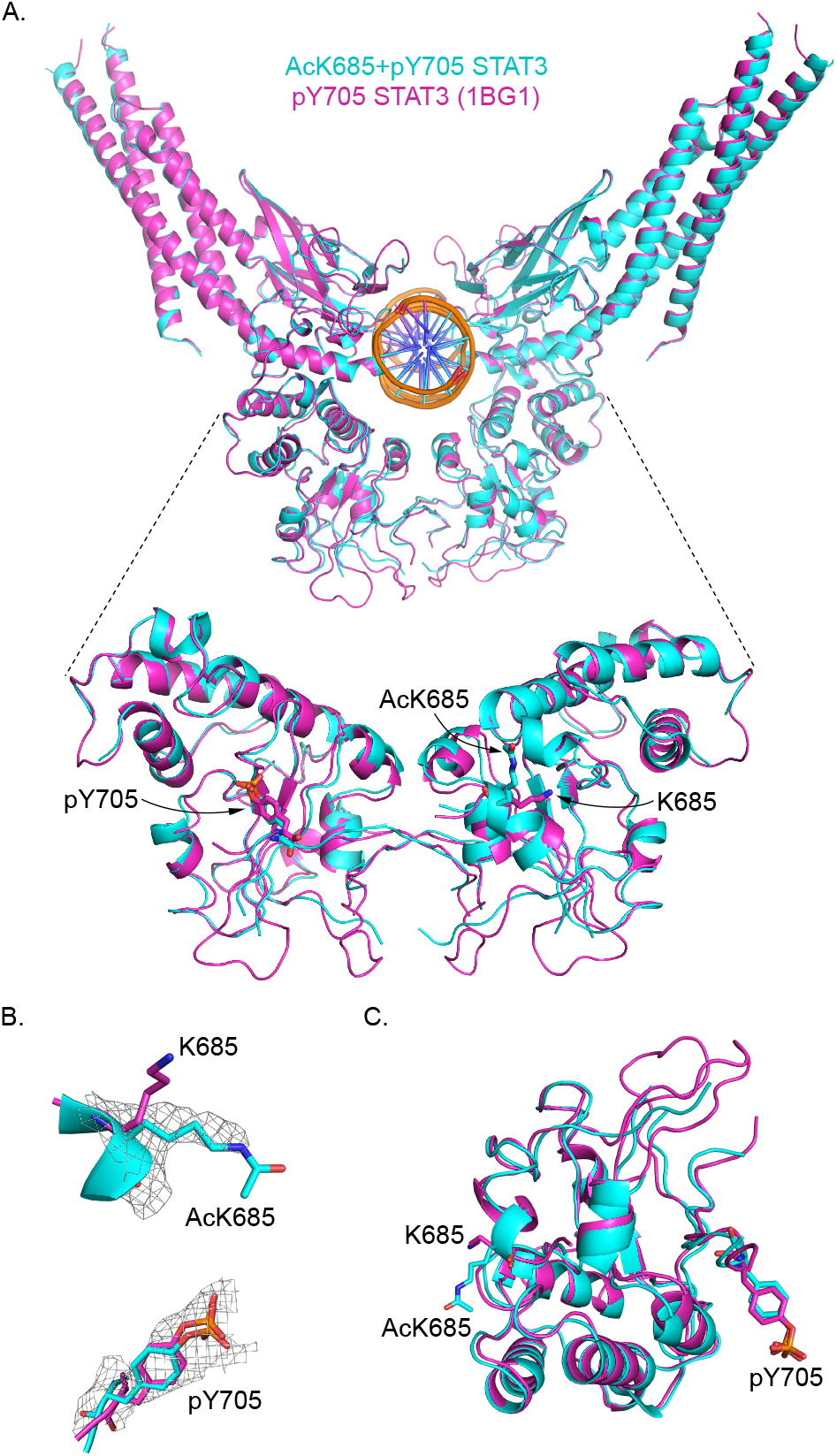
Crystal structure of AcK685+pY705 STAT3 in a complex with DNA. **A.** Superposition of AcK685+pY705 STAT3 (cyan) and pY705 STAT3 (magenta, PDB ID: 1BG1). Top: overall structure. Bottom: residues 500715, with residues AcK685, K685, and pY705 displayed in sticks model. **B.** Electron density map around residues AcK685 (top) and pY705 (bottom) (2*F_o_* – *F_c_* at 1.0*σ* and 0.7*σ* level, respectively), displayed relative to the position of the same residues in non-acetylated pY705 STAT3 (magenta). **C.** Position and orientation of residues AcK685, K685, and pY705 (in sticks model) within the SH2 domain of Lys685-acetylated and non-acetylated STAT3.

### 3.4. Deacetylation of Lys685-acetylated STAT3

An important aspect of any reversible post-translational modification, such as acetylation, is its regulation by enzymes that catalyze its removal. There are currently 18 known lysine deacetylases (KDACs) in the human genome, 7 nicotinamide adenine dinucleotide (NAD^+^)-dependent sirtuins (SIRT1–SIRT7) and 11 Zn^2+^-dependent histone deacetylases (HDAC1–HDAC11). To gain insight into the interaction between Lys685-acetylated STAT3 and mammalian KDACs, we used a semi-quantitative assay in bacteria to follow the deacetylation of AcK685 STAT3 and AcK685+pY705 STAT3 (Figure 4) [30]. In this assay, a KDAC is co-expressed in *E. coli* together with a C-terminal 6× His-tagged acetylated substrate, produced by genetically encoding the incorporation of an acetylated lysine. As such, the mammalian enzymatic deacetylation reaction is reconstituted in bacteria that serve as a ‘living test tube’. Deacetylase activity is then evaluated from Western blot analyses, by calculating the ratio between anti-acetyllysine immunoblot intensity (proportional to acetylation levels) and anti-6 × His immunoblot intensities (proportional to total protein levels). Using this methodology, we monitored the catalytic activity of SIRT1–7 and HDAC6 (residues 479-835, marked HDAC6*). We found that under these conditions, SIRT1–3 and HDAC6* were able to recognize AcK685 STAT3 and hydrolyze the acetyl group from position AcK685 (Figure 4, top panel). However, phosphorylation of position Tyr705 protected AcK685 STAT3 from deacetylation by SIRT1, SIRT3, and HDAC6*; AcK685+pY705 STAT3 could only be deacety-lated by SIRT2 (Figure 4, bottom panel). Hence, our data suggest that Tyr705 phosphorylation stabilizes the acetyl group on Lys685 by hindering the interactions with potential deacetylases.

**Figure 4:**
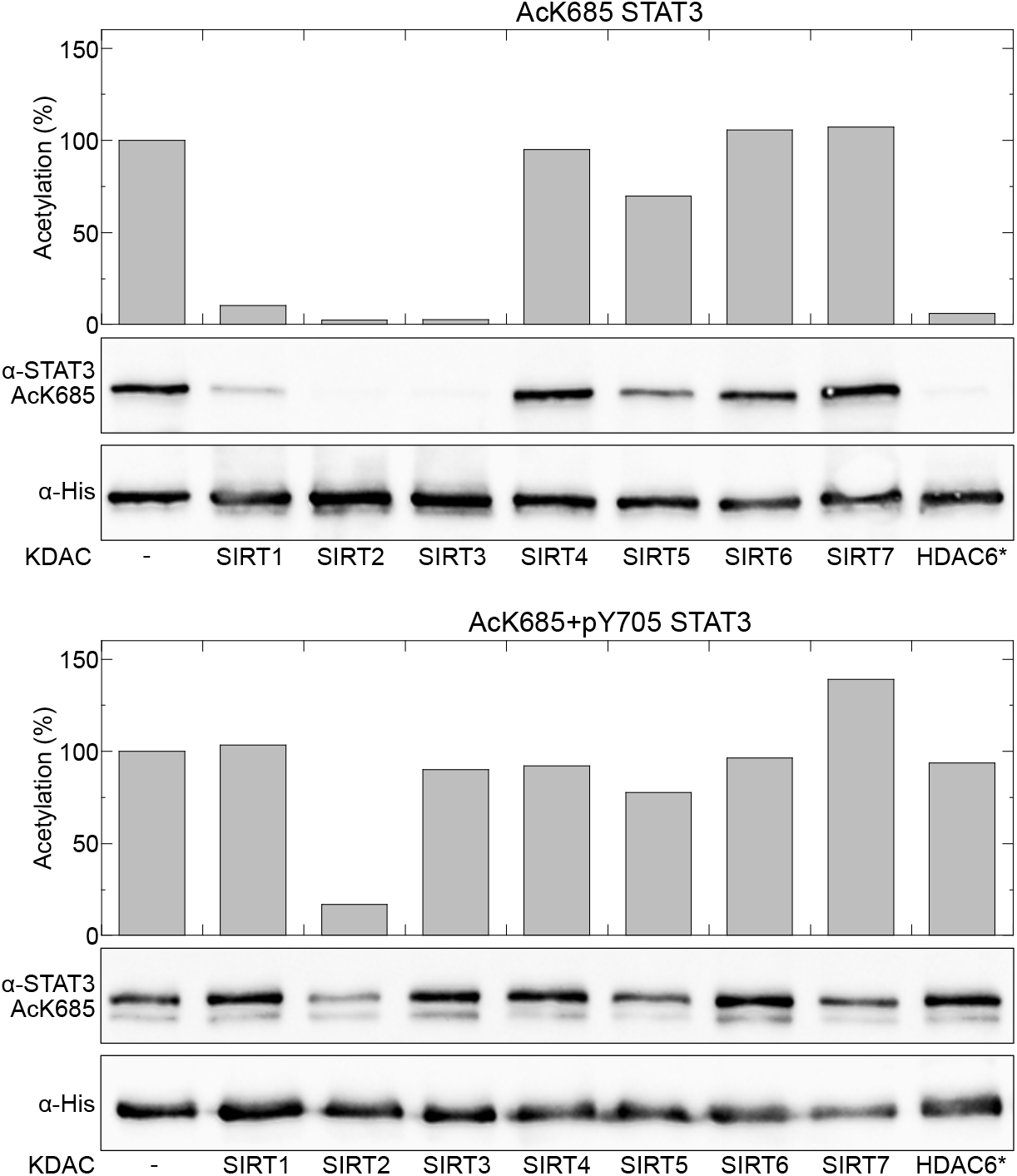
Deacetylation of STAT3 AcK685 in living bacteria. Indicated KDACs were co-expressed in bacteria with AcK685 STAT3 (top panel) or AcK685+pY705 STAT3 (bottom panel). Acetylation levels were calculated from the ratio between anti-STAT3 AcK685 and anti-6 × His immunoblot intensities and presented relative to the ratio calculated for the negative control (expression without KDAC). In this semi-quantitative assay, a given KDAC was considered active if the averaged ratio, calculated from at least three independent measurements, was less than 50% of the ratio calculated for the negative control. * HDAC6 was expressed as a truncated protein (residues 479–835), encompassing the second catalytic site.

## 4. Discussion

Previous in vivo studies demonstrated that acetylation of Lys685 promotes the dimerization and transcriptional activity of STAT3, suggestive of an acetylation-dependent mode of DNA binding and transcriptional activity. However, in the current study, we found no direct effect of Lys685 acetylation on STAT3 DNA-binding affinity or specificity. These results suggest that the acetylation-dependent STAT3 transcriptional activity observed in vivo may depend on other factors or conditions found in the complex cellular environment, such as additional post-translational modifications, protein-protein interactions, sub-cellular compartmentalization, etc. Alternatively, since the assays were performed with truncated proteins, the results may indicate that the N- and C-domains are important for acetylation-dependent transcriptional activity and protein-DNA interactions. Thus, our understanding of the role of Lys685 acetylation in STAT3 regulation could benefit from studies in cultured mammalian cells that utilize amber suppression technology to genetically encode the co-translational incorporation of an acetyl-lysine at position Lys685 of full-length STAT3. However, such studies are still technically challenging.

We found that Lys685 acetylation alone had no effect on the crystal structure of Tyr705-phosphorylated STAT3 in a complex with DNA. Several crystal structures of the STAT3 core domain have been determined, including the structure of non-phosphorylated STAT3 dimer in complex with DNA [3, 5]. These crystal structures and the structure presented here demonstrate essentially identical modes of DNA binding by STAT3. A possible explanation for the lack of any acetylation-dependent structural differences is that the crystal structure of STAT3 in complex with DNA represents only one DNA-binding mode, one which is not sensitive to Lys685 acetylation [40, 41].

Using deacetylation assay in bacteria, we found that SIRT1–3 and HDAC6* are capable of recognizing Lys685-acetylated STAT3 and catalyzing the hydrolysis of the acetyl group. Lys685-acetylated STAT3 is a known substrate of SIRT1, which predominantly resides in the nucleus and can also be found in cytoplasm [42–44]. HDAC6 shuttles between the nucleus and cytoplasm and was found to form a complex with the core domain of STAT3 [45]. SIRT3 is a major mitochondrial deacetylase, and STAT3 can translocate into mitochondria, where it regulates the mitochondrial respiratory chain via transcription-independent activity [9, 10, 46]. Interestingly, STAT3 residues Lys707 and Lys709 (but not Lys685) were found to be deacetylated by SIRT3 in mitochondria [11]. In addition, we found that deacetylation of AcK685 by SIRT1, SIRT3 and HDAC6* was dependent on the phosphorylation state of Tyr705. Considering that Tyr705-phosphorylation promotes STAT3 dimerization and that Lys685 is positioned close to the dimer interface, the observed phosphorylation-dependent protection might be explained by steric hindrance. This observation provokes the question of whether Lys685 acetylation promotes STAT3 dimerization and activation, or rather is the result of STAT3 Tyr705 phosphorylation-dependent dimerization. Dasgupta et al. found that Lys685 acetylation affects the transcriptional activity of un-phosphorylated STAT3 [19]. However, other studies found that Lys685 acetylation levels increase in response to stimulation by interferon-γ (IFN-γ), IFN-α, oncostatin M (OSM), interleukin 6 (IL-6), etc. As Tyr705 phosphorylation levels also increase in response to such treatments, one cannot rule out the option of phosphorylation-dependent increase in acetylation levels [16, 17, 20, 43]. Namely, Yuan et al. reported mutual increase in Tyr705 phosphorylation and Lys685 acetylation upon treatment with IFN-α and OSM [17]. Similarly, Wang et al. found increased levels of Lys685 acetylation in response to IL-6 stimulation [16]. Moreover, considering the observed effect of acetylation on STAT3 activity in vivo, phosphorylation of Tyr705 may activate STAT3 synergistically by promoting an increase in Lys685 acetylation levels.

Taken together, our data show no direct effect of Lys685 acetylation on the DNA-binding mode of STAT3. Numerous in vivo studies reported positive correlation between acetylation and transcriptional activity, dimerization, and nuclear translaocation of STAT3. Generally, in vivo studies are critical to our understanding of cellular processes, yet in vivo measurements of acetylation-dependent protein function are usually based on mutational analyses, gene knockdown or knockout, or use of deacetylase inhibitors. Consequently, any observed acetylation-dependent protein activity may be biased by indirect effects, such as altered histone modifications or transcription. Therefore, our work highlights the advantage of methodologies based on the site-specific incorporation of modified amino acids, as well as the importance of complementing in vivo studies with data obtained from in vitro measurements using homogeneously modified protein samples.

## Accession numbers

Coordinates and structure factors have been deposited in the Protein Data Bank with accession number 6QHD.

## Acknowledgments

The structural studies were performed on beamline ID29 at the European Synchrotron Radiation Facility (ESRF), Grenoble, France. We are grateful to the beamline scientists for providing assistance in using this beamline. We would also like to thank CCP4 staff for their contribution to the structure determination during the 1^st^ CCP4/BGU Structure Solution Workshop held at Ben-Gurion University in February, 2018. This work was funded by the Israel Science Foundation (grant number 807/15 to EA), the German-Israeli Foundation for Scientific Research and Development (grant number I-2342-203.13/2014 to EA) and by the U.S. National Institutes of Health (grant number 1R01GM117106 to RG).

