## Supporting information for "Unexpected implications of STAT3 acetylation revealed by genetic encoding of acetyl-lysine"

### Supplementary Tables

***Table S1.*** Data collection and refinement statistics.

| | |
|---|---|
| PDB code | 6QHD |
| Protein | AcK685+pY705 STAT3 |
| Beamline | ID29–ESRF |
| Wavelength (Å) | 0.97625 |
| Space group | $P4_1$ |
| **Cell Dimensions:** | |
| a, b, c (Å) | 175.49, 175.49, 79.07 |
| $\alpha$, $\beta$, $\gamma$ (°) | 90, 90, 90 |
| Resolution (Å) | 50–2.85 |
| Unique reflections | 47795 |
| Completeness (%) | 99.7 (98.4) |
| Mean $I/\sigma(I)$ | 10.22 (0.97) |
| Redundancy | 7.62 (7.81) |
| **Refinement:** | |
| $R_{work}$ | 0.294 |
| $R_{free}$ | 0.343 |
| $CC_{1/2}$ | 0.999 (0.653) |
| **Number of atoms (non hydrogens):** | |
| Protein | 9365 |
| Water | 89 |
| Average B-factor (Å$^2$) | 97.54 |
| Protein | 101.6 |
| Solvent | 81.59 |
| **R.M.S deviations:** | |
| Bond lengths (Å) | 0.019 |
| Bond angles (°) | 2.232 |
| **Ramachandran statistics:** | |
| Allowed | 349 (94.29%) |
| Partially allowed | 14 (4.07%) |
| Disallowed | 15 (1.65%) |

The values in parentheses refer to the data of the corresponding upper resolution shell. One crystal was used per data set. Data was collected at 100°K.
$R_{free}$ calculated using 5% of the reflection data chosen randomly and omitted from refinement.
R.M.S deviations for bonds and angles are the respective root-mean-square deviations from ideal values.

#### Supplementary Figures

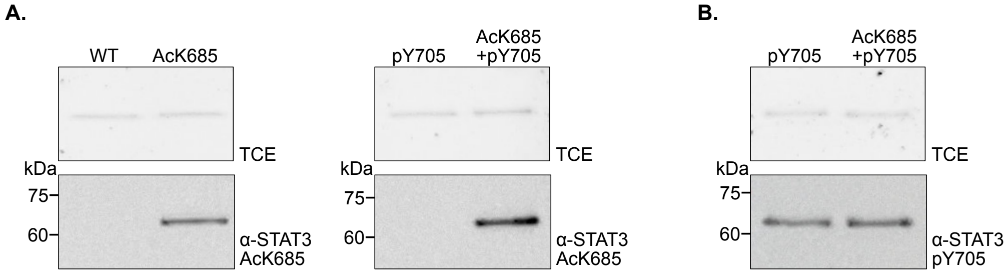


***Figure S1.*** Expression of STAT3 in bacteria. **A.** Indicated purified STAT3 proteins were analyzed by SDS-PAGE and visualized by Western blotting using specific antibodies against Lys685-acetylated STAT3. As loading control, proteins were also visualized by in-gel fluorescence following UV-induced reaction between tryptophan residues and 2,2,2-trichloroethanol (TCE). **B.** Indicated purified STAT3 proteins were analyzed as described in panel A, using specific antibodies against Tyr705-phosphorylated STAT3.

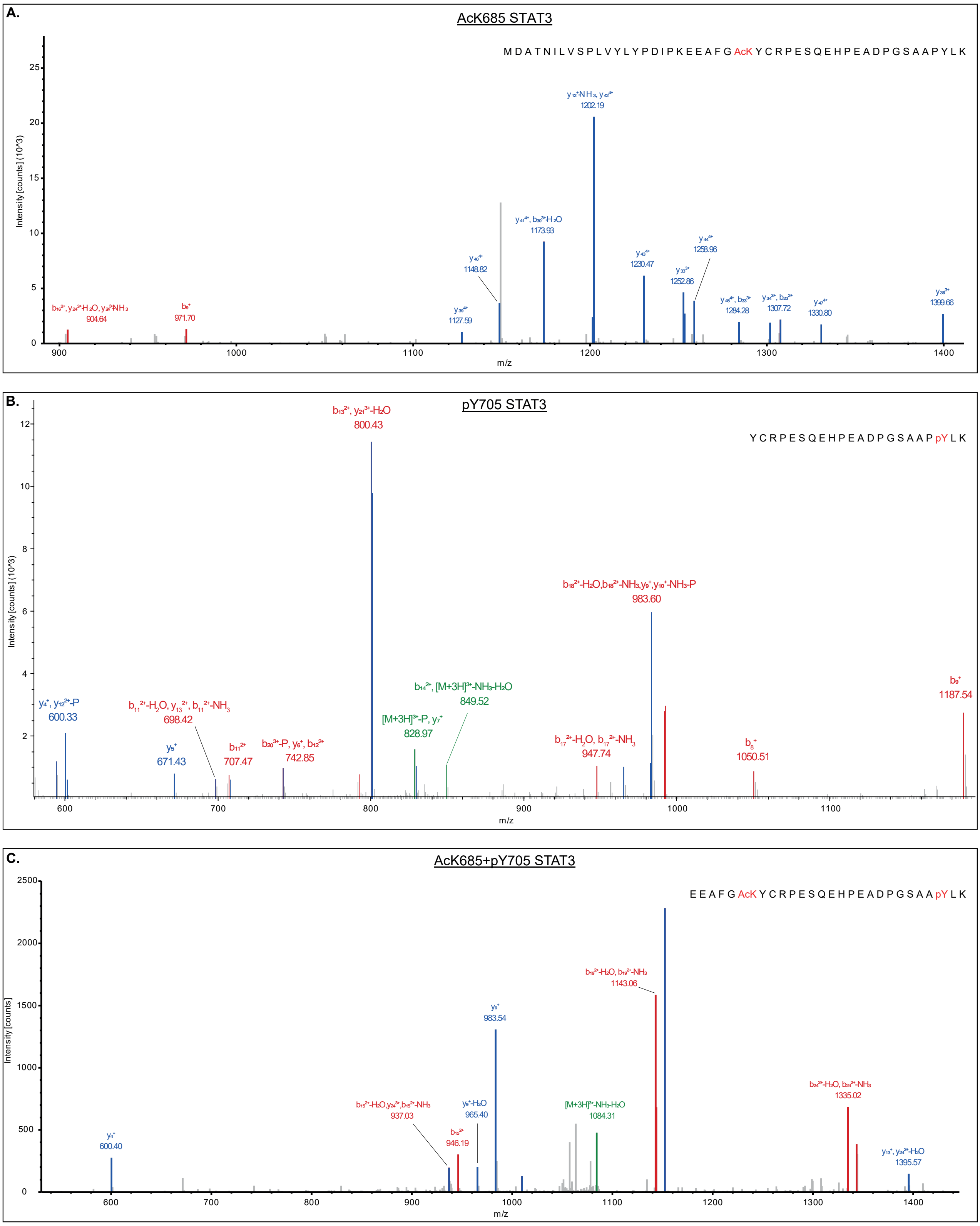


***Figure S2.*** LC-MS/MS identification of acetyl lysine and phosphotyrosine. MS/MS data confirm the incorporation of acetyl lysine at position 685 of AcK685 STAT3 (**A**), phosphotyrosine at position 705 of pY705 STAT3 (**B**), and both acetyl-lysine and phosphotyrosine at positions 685 and 705, respectively, of AcK685+pY705 STAT3 (**C**).

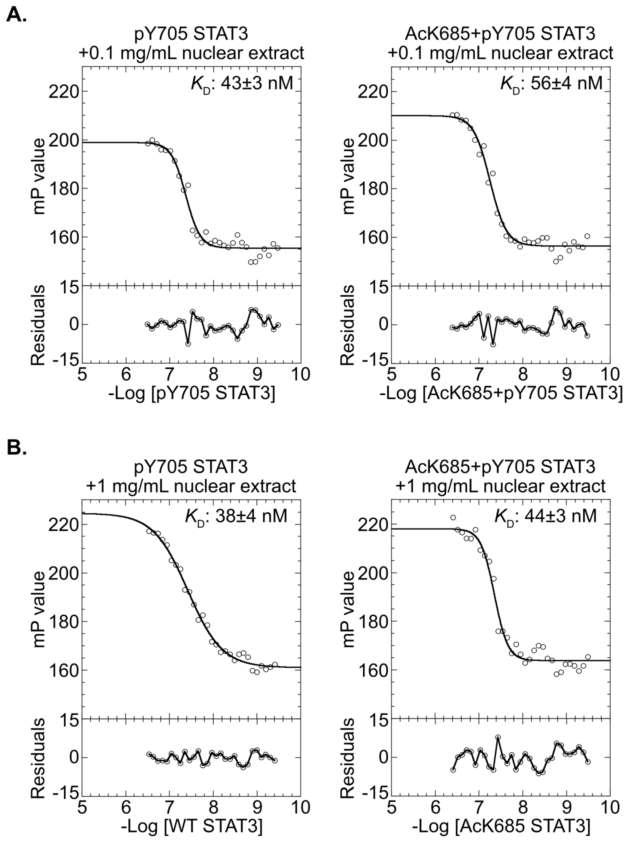


***Figure S3.*** Affinity of STAT3 variants to the binding site of the α2M promoter, was measured by fluorescence anisotropy in the presence of nuclear extract. Increasing concentrations of the indicated STAT3 proteins were incubated with 2 nM of 3'-fluorescein-labeled probe, and either 0.1 mg/mL (**A.**) or 1 mg/mL (**B.**) nuclear protein extract of STAT3-null cell line PC3. Data were fitted to Equation **??** and average $K_D$ ($n$=3, ±SD). Addition of nuclear extract was accompanied by an expected, yet not significant, increase in $K_D$ values. Importantly, the affinities of pY705 STAT3 and AcK685+pY705 STAT3 to α2M promoter were similar in the presence of 0.1 or 1 mg/mL nuclear extract. Thus, we were unable to isolate the cellular factor/s that interact differentially with pY705 STAT3 or AcK685+pY705 STAT3.

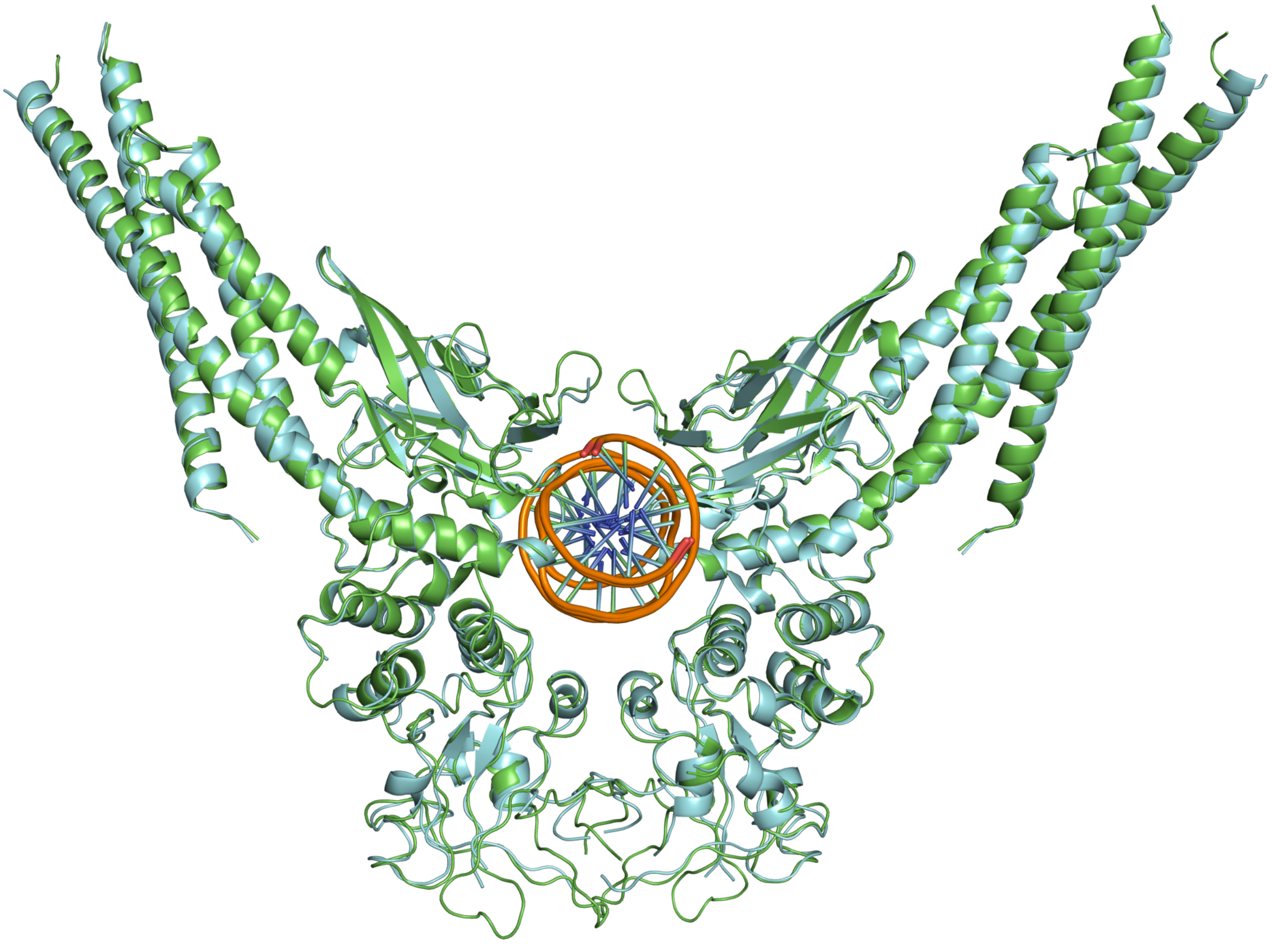

***Figure S4.*** Superposition of AcK685+pY705 STAT3 (cyan) and low-resolution crystal structure of AcK685 STAT3 (green). The low $C_{\alpha}$RMSD (0.64 Å) between the two structures, indicates that acetylation of Lys685 had no effect on the crystal structure of pY705 STAT3 in complex with DNA.
